# Protists Exhibit Stronger and More Recent Oceanic Genetic Structure than Archaeplastida and Metazoa

**DOI:** 10.1101/2025.06.02.657477

**Authors:** Rubén González-Miguéns, Alex Gàlvez-Morante, Guifré Torruella, Cédric Berney, Elena Casacuberta, Iñaki Ruiz-Trillo

**Author notes:** Rubén González-Miguéns.

## Abstract

The global distribution of biodiversity is shaped by a complex interplay of evolutionary history and ecological processes. While biogeographic patterns are well defined for animals and plants, the global distributions of protists remain unclear. A key question is whether protists follow the same broad biogeographic principles as macroscopic life. To address this, we compiled 88 marine COI metabarcoding studies performing population-genetic analyses across ocean basins. Our results reveal that most protist phyla exhibit pronounced genetic structure among oceans, a pattern exceeding that reported for Archaeplastida and Metazoa. This likely reflects recent, and potentially human-mediated, introductions, influencing protist dispersal and contemporary community assembly. By demonstrating that protist distributions are not historically cosmopolitan, our study supports the existence of common eukaryotic biogeographic patterns that transcend organismal size.

---

Unicellular eukaryotes (protists), which represent most of the diversity of eukaryotes from which multicellular groups evolved, have classically been viewed as fundamentally different from multicellular plants and animals with regard to their biogeographic patterns (*1*). Early theories postulated that protists have cosmopolitan distributions and virtually unlimited dispersal capacity, giving rise to the famous concept “everything is everywhere, but the environment selects” (*2*). This view, however, was based largely on limited morphological data. The advent of large-scale molecular techniques, such as metabarcoding, now permits better characterization of protist distributions (*3*).

Some recent marine metabarcoding studies have begun to reveal that protists exhibit biogeographic patterns more similar to those of multicellular eukaryotes than previously assumed (*4–6*). Nevertheless, these investigations have predominantly focused on presence-absence matrices, beta diversity patterns, and relative abundances inferred from read depth. A critical knowledge gap persists regarding population-level genetic diversity and the historical biogeographic processes shaping these distributions (*7*). This gap hinders our ability to determine whether the biogeographic principles observed in multicellular animals and plants are truly universal across all eukaryotes, or if smaller taxa follow distinct rules in marine environments. To resolve this metabarcoding offers a promising approach (*11*), as the large amount of molecular and geographic data from different organisms enables the integration of geographical and population genetic theories within a unified biogeographic framework (*8–10*). However, its effectiveness depends on meeting certain conditions: i) the use of molecular markers with intraspecific resolution (for example, the mitochondrial cytochrome oxidase subunit I gene, COI, which has proven invaluable in animal biogeography (*12*)); ii) the availability of a comprehensive global database for such markers that includes data from both protists and multicellular eukaryotes (*13*); iii) the use of standardized taxonomic units that mitigate the inherent biases of metabarcoding (*14*); and iv) the integration of population genetic methods and theory into metabarcoding analyses.

## Construction of a database with informative OTUs to test biogeographic patterns

To investigate global eukaryotic biogeographic patterns at the population level, a critical first step is the creation of a comprehensive metabarcoding database amenable to population-genetic analyses. So, we here compiled data from 88 independent metabarcoding studies into an environmental-COI (eKOI metabarcoding) database, comprising 976,865 amplicon-sequence variants (ASVs) with no abundance filter (Fig. S1) [see supplementary data (*15*)]; and 302,809,791 reads from 4,102 samples collected at 957 unique sites that span all oceans (Fig. 1a). We then taxonomically annotated them at phylum level, clustering them into 51,558 operational taxonomic units (OTUs).

**Fig. 1.**
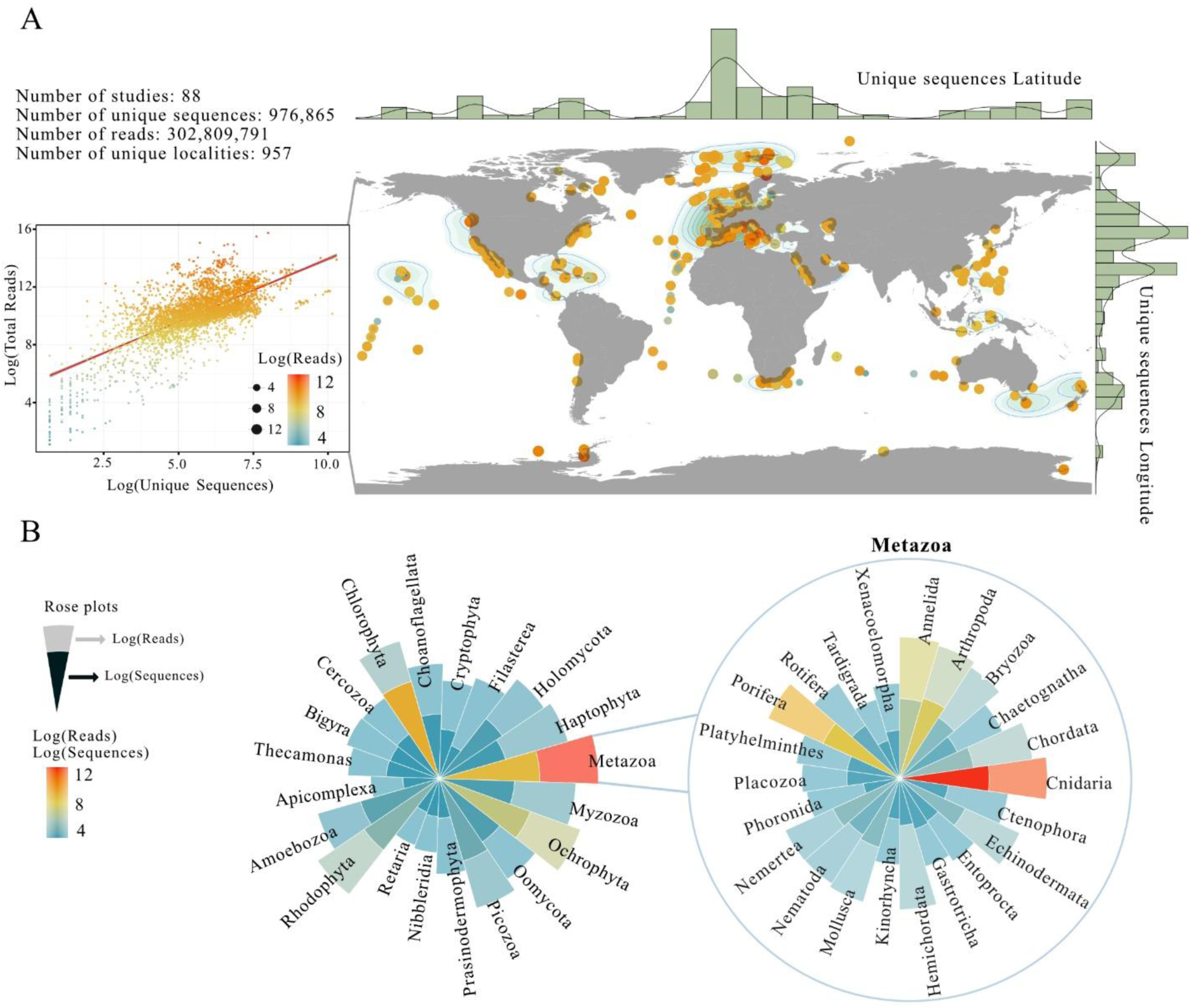
eKOI metabarcoding database. (**A**) Map showing the locations of samples included in the database. The size and color of the points represent the log(total reads) for each sample. To the left of the map, a scatter plot displays the relationship between log(ASVs) and log(reads) for the sampled localities, colored according to log(reads). (**B**) rose plots representing the proportion of taxonomic annotated ASV as log(ASV) on the inner layer and log(reads) on the outer layer in the different eukaryotic phyla used in the analysis.

To ensure robust population-genetic analyses and minimize potential artifacts, we stringently filtered the data to retain only informative OTUs, defined as those i) containing at least four sequences from a minimum of two distinct locations, and ii) containing at least one pair of sequences that differ by ≥ 1% (uncorrected p-distance). This rigorous filtering resulted in a final dataset of 5,970 informative OTUs, comprising 276,303 ASVs and 101,898,078 reads distributed across 41 phyla (Fig. 1b, 2a). Crucially, inter-phylum comparisons of nucleotide diversity (π) and Tajima’s D revealed minimal significant differences (0.001% and 6.9% of pairwise comparisons, respectively; Fig. S2; Supplementary Data), confirming broadly comparable molecular-evolutionary pressures across phyla. Consequently, this constructed eKOI metabarcoding resource provides standardized taxonomic units, thereby establishing a solid foundation for rigorous, population-level investigations of marine biogeographic hypotheses.

**Fig. 2.**
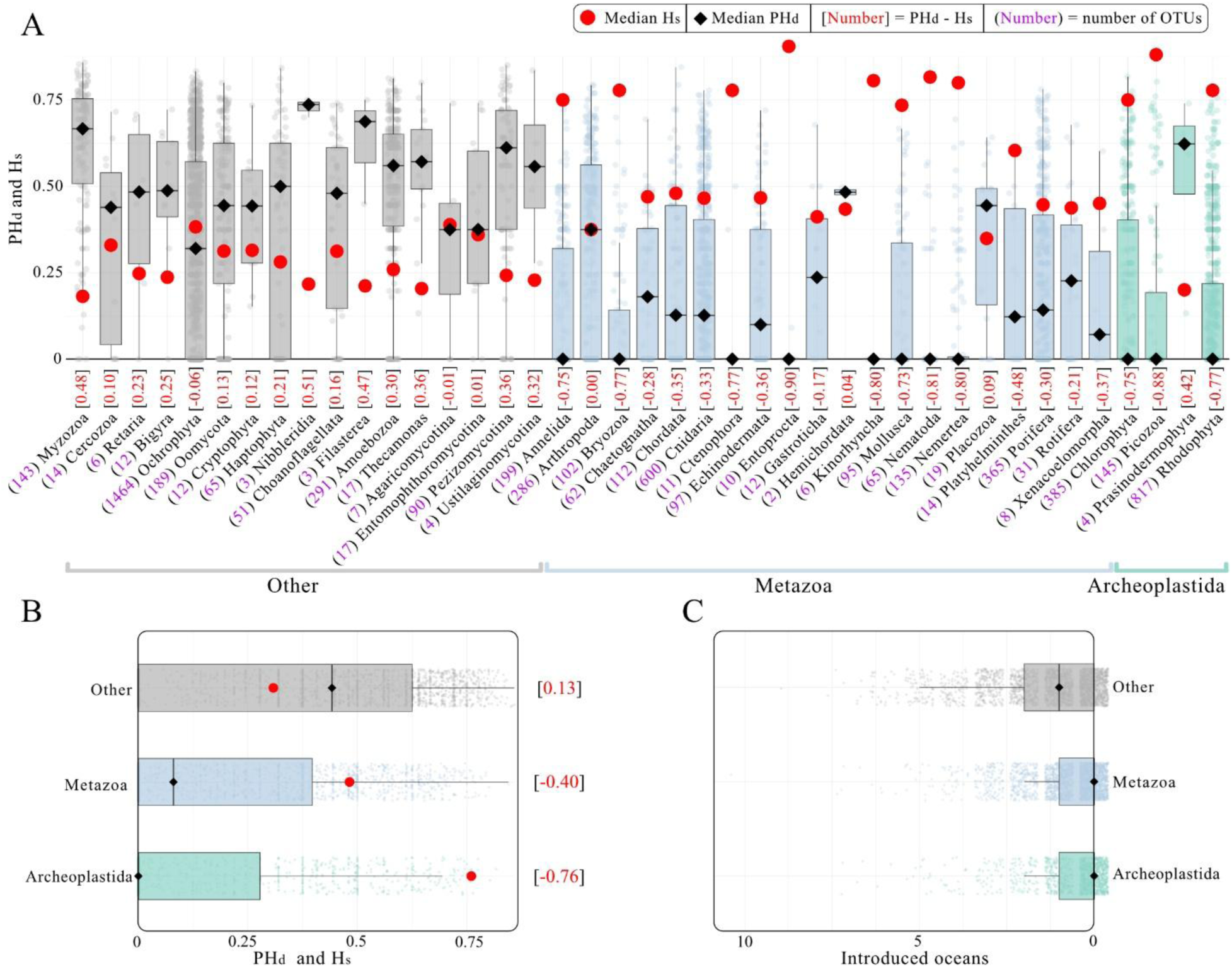
Population structure between eukaryotic phyla. (**A**) Boxplots representing the PH_d_ values for each OTU between phyla, where values close to 0 indicate low structuring in terms of intra-OTU haplotype diversity. Red circles with crosses represent the median H_s_ values for each phyla. The number near each phyla name represents the number of OTU recovered after filtration. The numbers at the top indicate the difference between PH_d_ and H_s_. (**B**) PH_d_ values grouped between different phyla. (**C**) Number of recent introduced oceans per OTU grouped between different phyla.

## Protists have larger geographic ranges than animals and plants

A long-standing debate in eukaryote biogeography concerns whether protists possess larger geographic ranges than macro-organisms (*1*). Testing this hypothesis has been hindered by the sheer volume of molecular and geographic species-level data required at a global scale (*16*). Our eKOI database allows for a direct assessment of this question. We analyzed the distributional patterns of all eukaryotic phyla by comparing the range size of every informative OTU within each eukaryotic phylum, quantifying i) the mean pairwise geographic distance among sampling sites, and ii) the maximum distance between any two sites. The results show that while Metazoa and Archaeplastida show mean inter-sample distances around 1,000 km, most protist phyla average over 5,000 km, with some individual OTUs exceeding 10,000 km (Fig. S3). This demonstrates that protists, at the OTU level defined by COI, possess considerably larger geographic ranges than plants and animals.

## Molecular and geographic expansions in protists

To understand why protists exhibit these distinctive geographic distances patterns, additional data, particularly molecular data, are essential, since they enable the detection of within-OTU genetic patterns linked to geographic range (*9*). Therefore, we compared genetic versus geographic distances within each informative OTU to test whether the observed spatial patterns were non-random and consistent across different phyla. Our analyses using both linear and nonlinear regressions revealed that most OTUs displayed significant non-random spatial patterns. Specifically, when compared to null models, 81.5% of OTUs in the linear models and 92.6% in the nonlinear models had R² values that were statistically significant (p < 0.05; Supplementary Data). Furthermore, linear and nonlinear models (Fig. S4), as well as Mantel tests (Fig. S5), consistently showed positive relationships between genetic and geographic distances across all eukaryotic phyla examined. In other words, genetic divergence tended to increase with greater geographic separation. The rate of genetic divergence per unit distance, quantified by the slope of the linear regression and by the parameter “a” of the nonlinear (exponential) model, was largely uniform across phyla. Significant differences among phyla in these rates were rare, occurring in only 5.7% of pairwise comparisons for the linear slopes and 3.6% for the nonlinear “a” parameters. Moreover, the geographic extent of an OTU (mean inter-sample distance) was only very weakly correlated with its genetic divergence rate, explaining merely 3.4% of the variance in slope and 4.0% in the “a” parameter. Together, these results indicate that geographic distance alone does not strongly limit genetic differentiation in protists, even across vast oceanic distances. We propose that the primary explanation for this weak isolation-by-distance pattern is the recent introduction and rapid geographic expansion of many protist species (*17*, *18*). Such rapid expansion over short evolutionary timescales would limit the accumulation of substantial genetic divergence, thus maintaining high genetic homogeneity across geographically distant populations. Nevertheless, this interpretation should be treated with caution, as multiple introductions can sometimes reduce genetic founder effects (*19*), yet molecular diversity often remains lower compared to that observed in the centers of origin.

## Metazoa exhibit higher richness of haplotypes among eukaryotes

To follow up on those rapid geographic expansions, we then inquired how and when current species distributions were established. To achieve this, we performed explicit population-genetic analyses to determine how molecular diversity is structured within each informative OTU. Our approach involved examining the genetic structure of these OTUs across different oceanic regions, treating each region as a distinct population. This allows for inferences regarding both contemporary and historical processes shaping current species distributions. To ensure unbiased analysis, population-level units within each OTU were defined irrespective of sequencing depth variations. We observed that haplotype richness per informative OTU had only a weak correlation with sequencing depth (R² = 18.2%), therefore being a robust metric for inferring population-genetic patterns in metabarcoding studies. Using the haplotypes identified within each OTU, we constructed haplotype networks for each informative OTU, without imposing a strictly bifurcating tree topology (*20*). This network-based approach allowed us to infer intra-OTU genealogies while avoiding the assumption of purely bifurcating relationships inherent to traditional phylogenies, which may not accurately capture genetic structure at the population level (*21*).

As a first step, we characterized haplotype diversity distributions across major phyla using three complementary metrics: haplotype diversity (H_d_) (*22*), branch diversity (B_d_), and a combined haplotype-branch diversity index (H_bd_) (*23*). These metrics were calculated based on the number of ASVs (unique haplotypes), the total read abundance per haplotype, and log-transformed read counts. B_d_ and H_bd_ values showed relative consistency across metazoan phyla, with few significant inter-phylum differences (Fig. S6). In contrast, H_d_ exhibited greater variation across phyla, primarily driven by comparisons involving Metazoa, which consistently displayed near-maximal H_d_ values. This indicates a significantly higher richness of unique haplotypes and, consequently, greater intra-OTU genetic diversity within metazoan OTUs compared to those of other eukaryotic phyla.

## Genetic Structuring Among Oceanic Regions suggest recent OTU introduction events in protist phyla

Following the characterization of within-phylum haplotype diversity, we next assessed genetic structuring across oceanic regions. For each informative OTU, we analysed haplotype frequency and distribution patterns across sampled oceans. We quantified population-level differentiation using four standard metrics of population genetic structure: total genetic diversity (H_t_), within-ocean genetic diversity (H_s_), the fixation index (F_st_) (a measure of among-ocean differentiation) (*24*), and the effective number of migrants (N_m_) (an indirect estimate of gene flow) (*25*). These metrics revealed significant differences among phyla in their degree of genetic structuring, with 20% of inter-phyla comparisons showing a significant difference in H_t_, 9.7% in H_s_, 21.9% in F_st_, and 9.3% in N_m_ (Fig. S7). The most pronounced disparities were observed in comparisons involving Metazoa and Archaeplastida versus other phyla. Specifically, Metazoa and Archaeplastida showed higher H_s_ values, reflecting greater average haplotypic diversity within individual oceans. This inter-oceanic diversity was coupled with very low F_st_ values in both clades (often approaching zero and never exceeding 0.5), signifying minimal genetic differentiation among oceanic regions. So, Metazoa and Archaeplastida haplotypes tend to be geographically restricted. In contrast, protist phyla exhibited much stronger genetic structuring, with consistently lower Hs and high F_st_ values (≥ 0.5). This pattern indicates that although lineages in those phyla occur across multiple oceans, their populations remain genetically isolated between regions. We propose that this strong structure in protists may reflect recent geographic expansions or introductions rather than long-term historical isolation.

To address the limitations of H_s_, in capturing haplotype distribution across oceans, we developed the Population-level Haplotype Diversity (PH_d_) metric, which integrates intra-oceanic diversity and the evenness of haplotype distribution across oceans [see supplementary materials and methods (*15*)]. PH_d_ penalizes uneven distributions, and the difference between PH_d_ and H_s_ (PH_d_ - H_s_) indicates the degree to which an OTUs genetic diversity is structured between oceans. PH_d_ analysis revealed significant inter-phylum differences, 20.6% of comparisons, largely mirroring H_s_ and F_st_ trends. Metazoa, Archaeplastida, and Ochrophyta showed near-zero PH_d_ values and negative PH_d_ - H_s_ differences, suggesting concentrated haplotypic diversity in limited oceanic regions (Fig 2a and b). However, some metazoan groups such as Arthropoda (PH_d_ - H_s_ = 0), Gastrotricha (-0.17), Placozoa (0.09), and Rotifera (-0.21), exhibited more even haplotype distributions. Protist phyla generally showed slightly higher PH_d_ values and PH_d_ - H_s_ differences close to zero, indicating moderate population genetic structuration with traceable dominant oceans. Overall, PH_d_ analysis supports the hypothesis of recent geographic expansions or introductions shaping genetic patterns in these marine eukaryotes.

Further analysis using the PH_d_ metric to identify dominant source oceans, harboring the majority of that OTU’s haplotypic diversity, and introduced (recipient) oceans revealed clear differences among taxonomic groups. Metazoa, Archaeplastida, and Ochrophyta showed minimal evidence of multi-ocean distributions (Fig. 2C, S8). In contrast, certain metazoan and protist phyla frequently displayed one or more introduced oceans per OTU, indicating recent inter-ocean colonization. For example, Arthropoda, Gastrotricha, Porifera, Rotifera, and Xenacoelomorpha, often had broader distributions, averaging around one introduced ocean per OTU (Fig. S8). Similarly, protists frequently displayed one or more introduced oceans per OTU, indicating a strong signature of recent inter-ocean introduction, with low haplotypic diversity, in these phyla. However, caution is required when characterizing species as native or introduced, as the evolutionary history of some species remains unclear with the current data (i.e, cryptogenic species) (*26*). To minimize misclassifications, we only considered OTUs with clear patterns of intra-OTU diversity across oceans [see supplementary materials and methods (*15*)].

To validate the capacity of our approach to identify genuine introduction events, we examined specific OTUs within the phylum Chordata, where introduction histories are often well-documented (*27*). Our analysis successfully highlighted several OTUs corresponding to species known for human-mediated introductions or widespread dispersal, such as *Acipenser gueldenstaedtii*, *Botryllus schlosseri*, *Salmo trutta*, *Siganus rivulatus*, and *Styela plicata* [see supplementary data (*15*)]. This reinforced the reliability of our approach in identifying recent introductions from metabarcoding data.

Furthermore, the inferred geographic scope of these colonization events differed markedly between groups. In Metazoa and Archaeplastida, most inferred source-sink connections occurred between geographically adjacent oceans and within the same hemisphere, suggesting predominantly regional spread (Fig. S9). In contrast, protist phyla OTUs often showed long-distance dispersal between non-adjacent oceans and different hemispheres, implying that recent introductions of these microorganisms frequently span distant oceanic regions, far exceeding the regional spread seen in larger eukaryotes (*28*). Finally, by aggregating dominant and introduced ocean assignments across all phyla, we uncovered broad biogeographic trends that align with known global patterns of species introductions [see supplementary materials and methods (*15*)]. The highest numbers of introduced informative OTUs were detected in Northern Hemisphere oceans, particularly the Atlantic and North Pacific (Fig. 3), consistent with these regions’ historically high rates of species introductions (*27*, *29*, *30*). In contrast, the main source regions (dominant oceans) across taxa were the Mediterranean Sea and Indian Ocean, both noted for their high levels of endemic diversity (*31*, *32*). Overall, these findings underscore that recent colonization, most likely facilitated by anthropogenic activities, have left a distinct imprint on global marine genetic diversity, with identifiable source areas and introduction hotspots explaining the distribution patterns we observe.

**Fig. 3.**
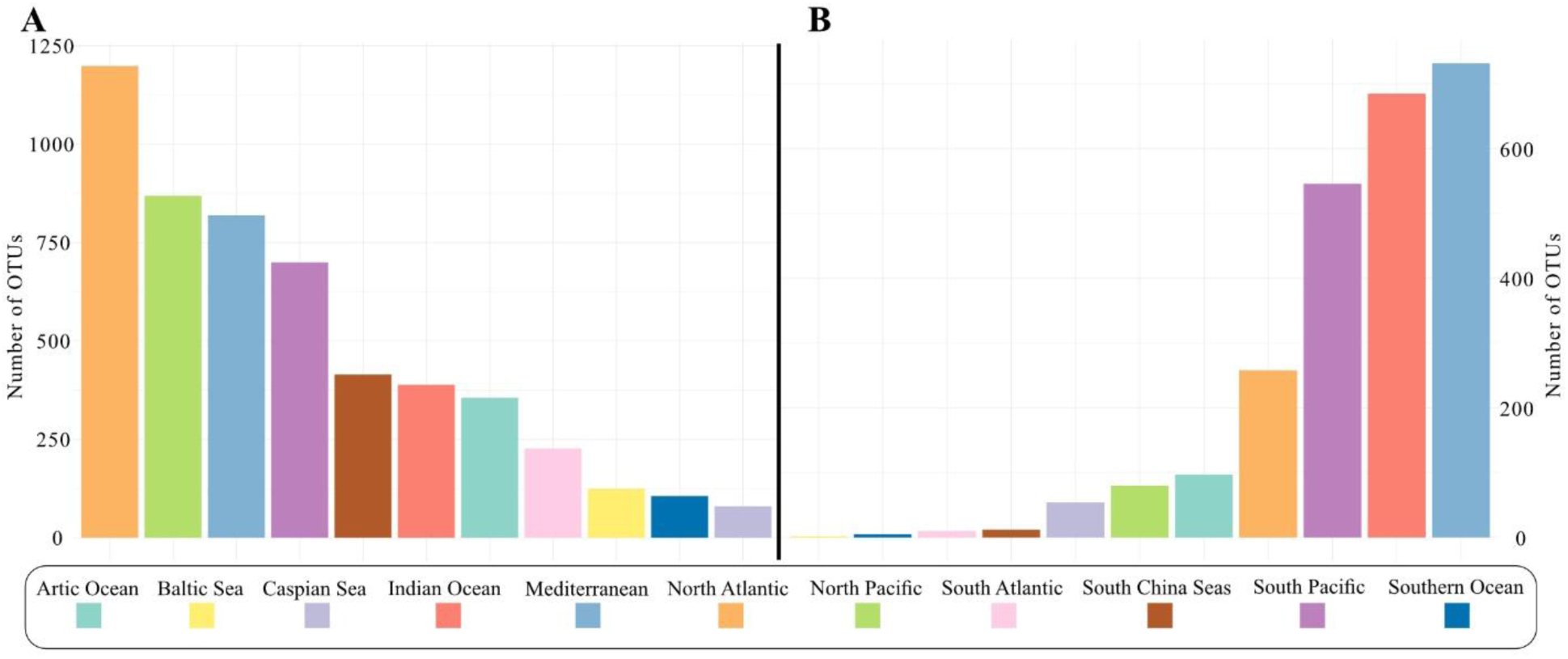
Number of recently introduced and dominant informative OTUs per ocean. (**A**) Boxplots showing the number of informative OTUs considered invasive in each ocean and (**B**) the number of informative OTUs per ocean identified as their dominant/sink.

## Conclusion

Our global COI analysis, leveraging informative OTUs and the innovative PHd metric for assessing population-level genetic diversity, definitively reveals fundamental distinctions in the contemporary biogeography of marine eukaryotes. Metazoa and Archaeplastida generally exhibit significantly restricted ranges and weaker ocean-basin genetic structure compared to protist phyla, underscoring divergent molecular population dynamics across major lineages.

These patterns likely reflect the profound impact of contemporary events, notably anthropogenic dispersal (*33*). While global shipping is a known vector for metazoan introductions (*34*), our data robustly indicate that protists and smaller metazoans (Arthropoda, Rotifera, Gastrotricha, Placozoa) show heightened susceptibility to long-distance dispersal, evidenced by elevated PH_d_ values with respect other metazoan phyla. Moreover, the lower PH_d_ in these smaller metazoans compared to protists phyla likely stems from protists rapid generation times (*35*), or multiple introductions (*19*), could facilitate the genetic homogenization in newly colonized oceans, reducing the founder effect. Conversely, restricted distributions and weak structure in some small and unicellular Archaeplastida suggest lower transoceanic transport survival, potentially due to light and resource limitations, corroborated by the differential recovery of taxa in ballast water studies. Consistent with this explanation, studies of ballast water frequently recover resilient photosynthetic taxa such as Bacillariophyta and dinoflagellates, which are capable of producing resistant forms (*36*), whereas groups like Chlorophyta appear underrepresented (*37*, *38*). This suggests that ballast water may be a less effective vector for long-distance dispersal in certain Archaeplastida lineages.

Ultimately, our global study establishes a unified framework for interpreting broad biogeographic and population genetic patterns across diverse eukaryotic phyla. The congruent patterns of marine geographic distribution and genetic structure within each group, strongly suggest shared historical processes, such as recent introductions, shaping their contemporary ranges. These findings challenge the traditional view of protists as ubiquitously dispersed entities (*1*), at the population genetic level, and reveal potentially universal principles governing biogeographic dynamics and community assembly, that transcend organismal size and complexity, particularly in the context of the Anthropocene.

## Materials and Methods

### eKOI metabarcoding database

To generate the eKOI metabarcoding database we selected 88 marine metabarcoding studies, with publicly accessible raw sequencing data, that had been generated on an Illumina paired-end platform using the cytochrome oxidase subunit I (COI) molecular marker, and employed the following primers: HCO (TAAACTTCAGGGTGACCAAAAAATCA) (*39*), LCO (GGTCAACAAATCATAAAGATATTGG) (*39*), jgHCO2198 (TAIACYTCIGGRTGICCRAARAAYCA) (*40*), mlCOIintF (GGWACWGGWTGAACWGTWTAYCCYCC) (*41*), and mlCOIintF-XT (GGWACWRGWTGRACWITITAYCCYCC) (*42*). After obtaining the raw data for each study, we processed each dataset separately according to the protocol outlined in (*43*). As a first step, primer trimming and demultiplexing were performed with Cutadapt (version 2.8) (*44*), when necessary. Next, the resulting reads for each sample in each metabarcoding study were processed independently using the DADA2 R package (*45*), with minor modifications applied depending on the specific metabarcoding study. Chimeric sequences were subsequently removed using the DADA2 *removeBimeraDenovo* function. After inferring the amplicon sequence variants (ASVs), we stored them in a FASTA file using the following header format: >sequence_ID merged_sample={ ‘LOCALITY_ID1’: reads, ‘LOCALITY_ID2’: reads, … }. A separate metadata file contained information associated with each locality ID (see **metadata.csv** in the supplementary data). For each sample in the metadata, we then assigned the corresponding ocean or sea. To achieve this, we downloaded shapefile datasets from the “Global Oceans and Seas” database (version 1, accessed 05/11/2024) and “Natural Earth” (accessed 05/11/2024). Using each locality’s latitude and longitude coordinates, we determined and recorded the respective ocean or sea for that sample. For subsequent biogeographic analyses, we utilised the broad ocean regions defined in the “Global Oceans and Seas” database, which groups the world into 11 major seas and oceans. This approach was chosen to avoid excessive subdivision of oceanic regions.

All FASTA files generated by each metabarcoding study, in the above format, were then merged into a single file (**eKOI_metabarcoding_database.fasta** in the supplementary data) using a custom Python script (1_fastas_combine.py). This script utilises the Biopython library (*46*) to identify identical ASVs across studies and combine their locality information. Each unique ASV was assigned an identifier in the format eKOIX (where X is a unique number for each ASV). Once the final merged FASTA file was assembled, we checked it for chimeric sequences using the VSEARCH *uchime_denovo* command (version 2.14.1) (*47*).

Taxonomic assignment of the ASVs was performed using the eKOI taxonomy database (*13*). For this purpose, we used the script (5_taxonomic_assignation.py) develop in that study. This script created a separate folder for each FASTA file in the working directory. Within each folder, an Excel file was generated containing the taxonomic assignment information for each ASV (**eKOI_metabarcoding_database_taxonomic_annotations.xlsx** in the supplementary data), obtained via the VSEARCH *usearch_global* command. ASVs with less than 84% similarity to any reference sequence were not considered further. Finally, we generated separate FASTA files for each taxonomic level of interest; in this case, we chose the phylum level. The resulting ASV sequences for each phylum can be downloaded from the “Phyla_data” folder in the supplementary data as **eKOI_database.fasta** file per phylum.

## Sequence Data Processing and Analysis

### OTU Clustering

Once FASTA files were generated for each phylum, they were analysed independently. First, all metadata for each ASV was extracted using the script 2_generate_abundances_unique.py. Biopython was used to extract each ASV’s ID and match it with the metadata file, producing a DataFrame containing the sequence ID, geographic coordinates, reads and ocean for each record (**abundances_unique.csv** in the supplementary data per phylum). Because some ASVs were found in multiple localities, duplicate entries were created for each locality where an ASV occurred by appending a suffix “_dupN” (where N is the duplicate number) to the ASV’s ID, yielding a unique identifier for each locality occurrence of that ASV. To avoid biases from multiple sequencing replicates per locality (common in metabarcoding studies), only one ASV entry per locality was retained (duplicate entries with identical ASV ID and coordinates were removed). After this filtering, a comprehensive FASTA file was compiled, including all ASV sequences (with each locality-specific duplicate as a separate entry). This file was aligned using MAFFT version 7.490 (*48*), with optimised parameters (--auto, --ep, --op, --maxiterate, --large) (**aligned_sequences_mafft.fasta** in the supplementary data per phylum). Operational Taxonomic Units (OTUs) were then delineated from the alignment using the script 3_generate_OTU.py. In particular, VSEARCH clustering at 97% identity was employed to group sequences into OTUs. We set the clustering threshold at 97 % identity (i.e., a 3 % divergence cut-off) because COI barcoding studies of Amoebozoa (*43*), Cercozoa (*49*) and diverse animal taxa report similar values (*12*). Although each lineage can display its own diversification rate, published barcoding gaps generally fall between 2 % and 3 %, so we applied a uniform 3 % threshold across all phyla. Representative ASV (centroids) for each OTU were saved, and a mapping of each ASV to its OTU was generated as a .uc file (**otus.uc** in the supplementary data per phylum). The .uc file was subsequently processed to produce a tabular mapping file (**otus_mapping.txt** in the supplementary data per phylum) that records the assignment of each ASV ID to its corresponding OTU.

### Molecular and geographic distance Matrix Construction and OTU Filtering

After obtaining the OTUs, the relationship between genetic and geographical distances among ASVs was analysed using the R script 4_1_generate_molecular_and_geographic_distances.R. We used the Biostrings R package to convert the phylum-level ASVs alignments into DNAbin objects, suitable for distance calculations. Genetic distance matrices were computed under the K80 model (Kimura 2-parameter) (*50*), with gaps (insertions/deletions) excluded using the *pairwise.deletion* method. In parallel, a matrix of geographical distances between sampling localities was calculated from the coordinate data using the *distVincentyEllipsoid* function of the geosphere R package. The genetic and geographical distance matrices were both converted to long format and merged into a single data frame, so that each row contained a pairwise genetic distance, the corresponding geographical distance, and the associated OTU (and phylum). To ensure robust analysis, we first filtered the data to include only OTUs represented by at least four independent ASVs. Next, we further filtered these OTUs, called the informative OTUs, to retain only those in which at least one pair of sequences had a genetic distance ≥ 0.01 and at least one pair had a geographical distance ≥ 1 m. The ratio of the number of total OTUs and filtered informative OTUs per phylum can be downloaded from **ratio_total_OTUs_informative_OTUs_by_phylum.csv** in the supplementary data, and the ID of the informative OTUs in the file **informative_OTUs.txt** of each phylum. This two-step filtering removed OTUs with very few sequences or with negligible variation, which could otherwise introduce noise. Metabarcoding studies that uses general primers lack targeted taxonomic resolution and often retrieve fragmented sequences or sequences from multiple species, potentially biasing interpretations (*14*). Filtering out OTUs with uniformly low molecular divergence could mitigates these issues. While this approach might also exclude some genuine low genetic diversity cases (for example, populations that have undergone bottlenecks or are in decline), it primarily reduces spurious OTUs arising from methodological biases, improving the reliability of biogeographic patterns interpretations.

The results of each analyses of each informative OTUs were compiled into the file **informative_OTUs_results.csv**, in the supplementary data. In this file, each row corresponds to a informative OTU and each column the results of the analysis or metrics, such as: **Phylum** (the taxonomic phylum of that OTU), **OTU** (the OTU identifier, unique within each phylum), **Max_Distance** (the maximum geographical distance, in metres, between any two localities), **Mean_Distance** (the mean geographical distance among all locality pairs), **Std_Dev_Distance** (standard deviation of geographical distances among localities) and **Count** (number of pairwise ASV comparisons, genetic/geographical distance pairs).

### Within-OTU Diversity and Tajima’s D

To quantify genetic diversity within each informative OTU, we calculated the nucleotide diversity (π). This was done using the previously generated alignments per phylum: the *nuc.div* function (*51*) was applied with *pairwise.deletion* = TRUE to compute π for each OTU using the script 9_calculate_nucleotide_diversity_by_otu.R. The resulting values were recorded in the **Nucleotide_Diversity_Pi** column of **informative_OTUs_results.csv**. We also calculated Tajima’s D for each informative OTU to investigate potential signals of selection or demographic history (e.g. recent expansion, bottleneck) using the script 10_calculate_tajimasD_by_otu.R. The *tajima.test* function (*51*) was applied to the alignment data for each informative OTU. In theory, a negative Tajima’s D indicates an excess of rare variants consistent with recent population expansion or purifying selection, whereas a positive Tajima’s D suggests a deficiency of rare variants, which may indicate a population bottleneck or balancing selection (*52*). Given our filtering criteria (which ensured multiple sequences per OTU and some level of genetic divergence), we expected Tajima’s D values to trend negative. Each OTU’s Tajima’s D value was recorded in a **Tajima_s_D** column, with its associated p-values recorded in **P_Value_Normal** (normal approximation) and **P_Value_Beta** (beta distribution approximation) columns. Subsequently, we examined whether nucleotide diversity and Tajima’s D varied significantly among phyla. (For Tajima’s D, this comparison was restricted to OTUs with a Beta p-value < 0.05) We first checked the distribution of each variable: the Shapiro-Wilk test indicated that the data were not normally distributed. Therefore, a non-parametric Kruskal-Wallis test (using the *kruskal.test* function in R) was employed to test for differences among phyla. A significant Kruskal-Wallis p-value was taken as evidence of variation among phyla for the given metric. In cases of significance, post-hoc pairwise comparisons were performed using Dunn’s test (*dunnTest* function in R) with a Bonferroni adjustment for multiple testing (*53*). The results of the pairwise comparisons for each variable were saved to **significant_group_comparisons_Dunn_test.csv** (available in the Supplementary Data). The file contains the columns **Z** (Z-Statistic), **P_unadj** (Uncorrected p-value) and **P_adj** (Adjusted p-value) for every variable in the phyla pairwise comparison. Phyla with an insufficient number of informative OTUs were excluded from these comparative analyses (Filasterea, Hemichordata, Nibbleridia, Prasinodermophyta and Ustilaginomycotina). Additionally, for infer differences between and within groups, phyla were grouped into three broad categories (Archaeplastida, Metazoa, and all other phyla) to compare the prevalence of significant differences between and within these groups (**significant_group_comparations_Dunn_test.csv** in the supplementary data). The same statistical approach was applied in later analyses for other variables and metrics of interest, as described below.

### Read Count Filtering Impact

It is common in metabarcoding studies to discard ASVs with very low read counts (below a certain threshold) to avoid spurious ASVs (*54*). To evaluate how such filtering might affect the recovery of low-abundance haplotypes and OTUs, we characterised the occurrence of rare haplotypes within each OTU. First, for each informative OTU, the total number of ASVs and the total number of reads was recorded in the **Num_Sequences** and **Total_Reads** columns, respectively in the **informative_OTUs_results.csv** file. In addition, separate columns captured the number of reads contributed by each ocean named **<ocean>_reads**. We then evaluate the potential loss of data and in reads counts filtering by counting the number of haplotypes with fewer than 10 reads for each informative OTU, recorded in the **haplotypes_lt10** column. We also determined how many of these rare haplotypes were supported by at least two independent ASVs (each with <10 reads); this number was recorded in **haplotypes_recovered_multiple** column. By requiring multiple independent ASVs, we can identify rare haplotypes that are consistently observed, thereby reducing the likelihood of counting sequencing artifacts as true haplotypes. We also assessed whether the low-read filter could inadvertently affect more abundant haplotypes. For each OTU, we counted the number of haplotypes with total reads > 10 that nevertheless contained at least one constituent ASV with <10 reads. This count was recorded in **haplotypes_gt10_with_lt10_seq** column. Finally, to examine the biogeographic distribution of low-abundance ASVs, we counted how many oceans had a total read count < 10 for each OTU. This value (the number of distinct oceans in which the OTU is represented by fewer than 10 reads) was recorded in the **global_lt10** column. Given that metabarcoding studies are typically constrained to specific regions/oceans, any potential “tag jumping” across oceans (a form of cross-sample contamination) can be ruled out at this stage. Overall, these metrics allowed us to understand the extent to which stringent read filtering might exclude genuine but low-frequency haplotypes and the veracity of ASVs with low number of reads.

## Geographic and genetic distances

Linear regression and non-linear analyses were conducted for each informative OTU to evaluate the relationship, strength and statistical significance of the correlation between genetic and geographic distances using the R scripts 5_calculate_linear_models_by_otu.R and 6_calculate_nls_models_by_otu.R, respectively. The parameters of the linear model (lm) were calculated for each informative OTU using the dplyr R package, and the results were saved in the file **informative_OTUs_results.csv**, with the Columns: **Intercept_lm** (intercept), S**lope_lm** (slope), **R2_lm** (coefficient of determination), **P_value _lm** (p-value of the slope), **AIC_lm** (Akaike Information Criterion), and **BIC_lm** (Bayesian Information Criterion). Additionally, a non-linear regression analysis (nls) was performed using the model: “Genetic Distance” = a * log(“Geographical Distance” + 1) + b. Parameters a and b were estimated from predefined initial values. The output included the **a_nls** (slope parameter), **b_nls** (vertical offset parameter), **RSS_nls** (residual sum of squares), **R2_nls** (coefficient of determination), **AIC_nls** (Akaike Information Criterion), and **BIC_nls** (Bayesian Information Criterion). To interpret the differences in AIC and BIC between these two models, categories were established based on the magnitude of the differences (Δ): (1) no evidence to prefer either model (|Δ| ≤ 2), indicating similar fit; (2) moderate evidence (2 < |Δ| ≤ 7), indicating moderate support for one model; and (3) strong evidence (|Δ| > 7), indicating strong support for one model (**models_comparison_summary.csv** file in supplementary data). Based on these results we continued with the non-linear models for the graphical representations, as it better explained the genetic and molecular distance correlations per phylum.

Next, the linear and non-linear models were compared with null models to assess whether the relationship between genetic and geographic distances was consistent with a stochastic pattern. The null models were generated by simulating random distributions of genetic and geographic distances using the Python script 7_calculate_null_models_by_otu.py, fitting linear models with the scikit-learn library and optimising non-linear models with SciPy’s curve_fit function. For each OTU, 1,000 simulations were performed. In each simulation, the number of data points equalled the number of pairwise comparisons (**count** column in **informative_OTUs_results.csv** file) for that OTU. Geographical distances were sampled uniformly between 0 and the maximum recorded distance (**max_distance** column), and genetic distances were sampled uniformly between 0 and 0.03 (the specified OTU cutoff). From these simulations, the mean, median and standard deviation of R^2, AIC and BIC were calculated for comparison with the observed models. Results were saved in **informative_OTUs_results.csv** file with the following columns: **Mean_R2_lm_null**, **Median_R2_lm_null**, **Std_R2_lm_null**, **Mean_AIC_lm_null**, **Median_AIC_lm_null**, **Std_AIC_lm_null**, **Mean_BIC_lm_null**, **Median_BIC_lm_null**, and **Std_BIC_lm_null**; and for non-linear models: **Mean_R2_nls_null**, **Median_R2_nls_null**, **Std_R2_nls_null**, **Mean_AIC_nls_null**, **Median_AIC_nls_null**, **Std_AIC_nls_null**, **Mean_BIC_nls_null**, **Median_BIC_nls_null**, and **Std_BIC_nls_null**, **Mean_a_nls_null**, **Median_a_nls_null**, **Std_a_nls_null**, **Mean_b_nls_null**, **Median_b_nls_null**, and **Std_b_nls_null**. P-values were calculated to determine whether the observed relationships differed significantly from random expectations, based on the coefficient of determination, and were saved in the columns **P_R2_lm_null** (linear models) and **P_R2_nls_null** (non-linear models).

To further examine whether a correlation exists between genetic and geographic distances and to assess its consistency across phyla, Mantel tests were performed using the R script 8_calculate_mantel_test_by_otu.R. For each OTU, genetic and geographic distance matrices were generated, and the Mantel test was conducted using the *mantel* function from the vegan package (version 2.6) (*55*) with 999 permutations, the results were recovered in the columns **Mantel_R** and **P_Value_Mantel**. However, the Mantel test can be problematic due to inherent dependencies in the data (*56*). In this context, the permutations used to calculate the p-value may not adequately reflect the expected null distribution under true independence; therefore, this results should be interpreted with caution. After completing these analyses, only OTUs with a Mantel test p-value < 0.05 were retained, and comparisons between phyla were conducted as described above.

## Biogeography Based on Haplotype Networks

### Haplotype network construction and diversity metrics

Unique haplotypes were first identified from the alignment for each informative OTU using the *haplotype* function (*51*). The number of haplotypes per informative OTU was then recorded in a column named **num_haplotypes**. Next, using the *haploNet* function (*51*), haplotype networks based on Minimum Spanning Networks (MSNs) were constructed for each informative OTU and used in subsequent analyses. We then calculate the Haplotype diversity (H_d_), branch diversity (B_d_) and their combined metric (H_bd_) (*23*) using R script 11_calculate_haplotype_diversity_by_otu.R. We saved the results in columns named **Hd**, **Bd** and **HBd**, respectively, in the **informative_OTUs_results.csv** file. Additionally, the maximum value of the squared haplotype frequencies per OTU was recorded in a column **Efh2_Hd**.

The above metrics treat each ASV equally, regardless of its abundance (i.e. they are based only on ASV presence/absence within each haplotype). To account for the relative abundance of each ASV (as a proxy for its prevalence in the sample), we incorporated the associated read counts. Specifically, we used two metrics (1) the total reads count for each ASV and (2) a logarithmic transformation, (log(reads+1)), using the R script 12_calculate_weighted_haplotype_diversity_by_otu.R. By weighting sequences according to their total read counts and a logarithmic transformation, haplotypes with higher abundance have a greater influence on the calculated diversity metrics. Therefore, if a sequence-based diversity metric exceeds its read-weighted counterpart, this indicates that rarer haplotypes disproportionately influence the haplotypic diversity (i.e. diversity is overestimated in the unweighted case). This approach is particularly useful for communities with high variability in abundances, as it allows patterns of dominance or inequality to emerge. However, it may underestimate diversity in OTUs that contain many low-abundance haplotypes. The results of these abundance-weighted calculations were recorded in the following columns: **Hd_reads** and **Hd_log_reads** (haplotype diversity); **Bd_reads** and **Bd_log_reads** (branch diversity); **Hbd_reads** and **Hbd_log_reads** (combined haplotype and branch diversity); and **Efh2_reads** and **Efh2_log_reads** (maximum squared haplotype frequencies).

### Oceanic genetic structure

After calculating haplotype diversity metrics, we evaluated genetic structuring between and within populations (each corresponding to a different ocean). Analysis of Molecular Variance (AMOVA) was performed for each OTU using R script 13_calculate_amova_by_otu.R. OTUs were selected for AMOVA based on the following criteria: (i) Presence in at least two different oceans; (ii) At least two populations; and (iii) At least two distinct haplotypes within those populations. We used the *popr.amova* function to perform the AMOVA. Variance components and sums of squares were calculated for between-population and within-population comparisons. The results were saved in the **informative_OTUs_results.csv** file, in columns named **Between_samples_SumSq**, **Within_samples_SumSq**, **Total_SumSq**, **Between_samples_Sigma**, **Within_samples_Sigma** and **Total_Sigma**. These metrics were then compared across phyla as described above. Because the sequence length is relatively short (approximately 313 bp), the AMOVA results should be interpreted with caution.

To mitigate potential biases introduced by short sequences, we evaluated genetic structuring among oceans using the haplotype frequencies and distributions for each informative OTU. The haplotypic diversity metrics total diversity (H_t_), within-population diversity (H_s_), genetic differentiation coefficient (F_st_) and gene flow (N_m_) were calculated for each OTU using script 14_calculate_Hs_fst_nm_by_otu.R. These four metrics were stored in columns **Ht**, **Hs**, **Fst**, and **Nm**; and comparisons between phyla were conducted as described above.

A modified haplotype diversity metric (PH_d_) was created to infer the distribution of haplotypes among populations (here defined by different oceans) using script 15_calculate_PHd_and_global_ocean_dominance_by_otu.R. This new metric incorporates the relative distribution of haplotypes across populations. By using weighted relative haplotype frequencies, PH_d_ quantifies the contribution of population-level differences to the genetic diversity of each OTU. The classical haplotype diversity formula was adapted to account for the distribution of haplotypes across populations. In this adaptation, the relative frequency of each haplotype is redefined as its frequency in a given population divided by its total frequency in the OTU. The modified PH_d_ is therefore calculated as:

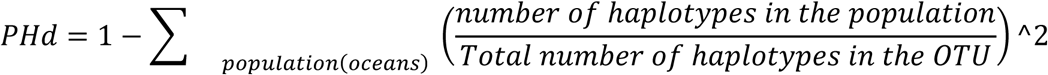

The PH_d_ values were saved in a column named **PHd**. The relative haplotype frequencies for each ocean were stored in columns named **<ocean>_sqfreq**. These PH_d_ metrics were then compared across phyla using the same procedure as for the other diversity metrics. Furthemore we compare PH_d_ with H_s_ metric to helps to identify whether the populations of an informative OTU are genetically structured or relatively homogeneous across oceans. If PH_d_ exceeds H_s_ (a positive difference), this indicates that the populations have distinct genetic compositions and are partially isolated from one another, reflecting significant genetic structuring and no single population dominating the haplotype diversity. Conversely, if PH_d_ is less than H_s_ (a negative difference), one or a few populations account for most of the global haplotypic diversity in the OTU, suggesting that these populations dominate the haplotypic diversity.

For each informative OTU, we identified the ocean with the highest haplotype relative contribution (the maximum sqfreq value) as a proxy for the putative population of source and vice versa for recent introduction populations. However, this analysis is considered a preliminary proxy to guide further investigation. To distinguish likely source populations from recent introductions, we calculated ratios of relative haplotype frequencies between oceans using script 20_introductions_vs_dominant_oceans_byOTU.R. Specifically, within each OTU we used the ocean with the highest relative frequency as the reference (ratio = 1), and we computed for each other ocean the ratio of the reference frequency to that ocean’s frequency. We then examined the distribution of these ratios across all informative OTUs to detect shifts toward lower values. A notable change in the distribution occurred at a ratio of 0.25. Consequently, for each OTU we classified any ocean with a ratio below 0.25 as a “recently introduced ocean/population,” since such oceans have very low relative haplotype frequency compared to the highest haplotypic frequency ocean. Furthermore, to identify a clear source ocean for each informative OTU, we applied additional filtering criteria. We selected OTUs in which exactly one ocean had a ratio of 1.0 (i.e. a single ocean had the maximum relative frequency) and excluded informative OTUs where multiple oceans shared this maximum frequency, since that would preclude a clear source. We also excluded informative OTUs in which any other ocean had a ratio exceeding 0.5, to avoid ambiguity. Although these thresholds are somewhat arbitrary, they provide a more restrictive criterion that allows a more reliable inference of the putative source ocean for the remaining informative OTUs. We create a pie chart graph of the proportion of dominant oceans per phylum **ancestral_oceans.pdf**. Furthermore, we combine the number of OTUs dominant and introduced oceans in the file **dominant_introduced_oceans.csv**.

To validate this proxy for source (dominant) and introduced ocean populations, we used phylum Chordata taxonomically annotated informative OTUs as a positive control. We included only those informative OTUs that had at least one ocean classified as “recently introduced”. For each selected informative OTU, the first amplicon sequence variant (ASV) was compared to entries in the NCBI database using BLAST (*57*). We retained those OTUs for which the BLAST hit showed ≥ 97% sequence identity to a species in the database, ensuring that species-level identification was feasible. Taxonomic annotation results for these OTUs are recorded in file **introduced_chordata_otus.csv**.

For each informative OTU, we inferred connectivity between different oceans from its haplotype network to evaluate genetic dispersal and exchange. This analysis was performed using the script 16_generate_ocean_connectivity_by_otu.R. In each haplotype network, nodes represent haplotypes and edges represent connections between them. We annotated each edge with the ocean of origin for each haplotype in the pair. Only connections linking haplotypes from different oceans were retained; intra-oceanic connections were discarded because they do not inform inter-ocean connectivity. We then aggregated the data to count the number of connections between each pair of oceans. For each informative OTU, we recorded the total number of oceanic haplotypic connections in the column **Total_Connections** and the number of connections per ocean in columns named **<ocean>_connections**. Then, we classified oceanic pairs as “nearby” if they are geographically adjacent and “distant” if they are geographically separated using the script 17_calculate_ocean_conectivity.R, the number of connections were recovered in the columns **nearby_oceans_connections** and **distant_oceans_connections**. OTUs in which all haplotypes occur in the same ocean therefore have only “nearby” connectivity and no “distant” connectivity. We also characterised intra-hemispheric and inter-hemispheric connectivity, focusing on connections between oceans in the Northern and Southern Hemispheres, recovered in the columns **Inter-Hemisferica** and **Intra-Hemisferica**.

For each phylum, we generated network graphs illustrating connections among oceans using the script 18_oceanic_network_by_phyla.py and 19_global_oceanic_network.py. In these graphs, each node represents an ocean and each edge represents the total number of connections between two oceans, weighted by relative strength. We summed all connections across OTUs for each phylum and applied a log1p transformation (the natural logarithm of one plus this sum) to normalise the data. We then applied the Louvain community detection algorithm to identify clusters (communities) within each network, colouring nodes by their community membership. The networks were constructed using the networkx library, with edge weights reflecting connection strength the network per phylum were recovered as **network_<phylum>.pdf and .jpg**.

Finally, we analysed correlations among the different calculated variables for each informative OTU. We computed correlation matrices and associated p-values using Pearson’s correlation coefficient, generating a table listing only those correlations that were statistically significant (p < 0.05) **significant_correlations.csv** file.

## Acknowledgments

we express our gratitude to A. Berlinches, E. Lara, G. Bercedo and M. Villar-DePablo and the MCG lab group for helpful discussions. Work funded by the European Union (ERC, MISSINGRELATIVES, 101097659). Views and opinions expressed are however those of the author(s) only and do not necessarily reflect those of the European Union or the European Research Council. Neither the European Union nor the granting authority can be held responsible for them. We also acknowledge support to Departament de Recerca i Universitats de la Generalitat de Catalunya (exp. 2021 SGR 00751) and support by PIE-202120E047-Conexiones-Life, and MICIU/AEI /10.13039/501100011033 - FSE+ (JDC2023-050439-I).

## Author contributions

Conceptualization: RGM, IRT

Methodology: RGM

Investigation: RGM

Visualization: RGM

Funding acquisition: RGM, IRT

Project administration: RGM

Supervision: EC, IRT

Database maintenance: AGM, GT, CB

Writing – original draft: RGM, IRT

Writing – review & editing: RGM, AGM, GT, CB, EC, IRT

## Data and materials availability

All data, code, results, including eKOI metabarcoding database, are available via Figshare (10.6084/m9.figshare.29144645). All other data are available in the manuscript or the supplementary materials.

---

**Fig. S1.**
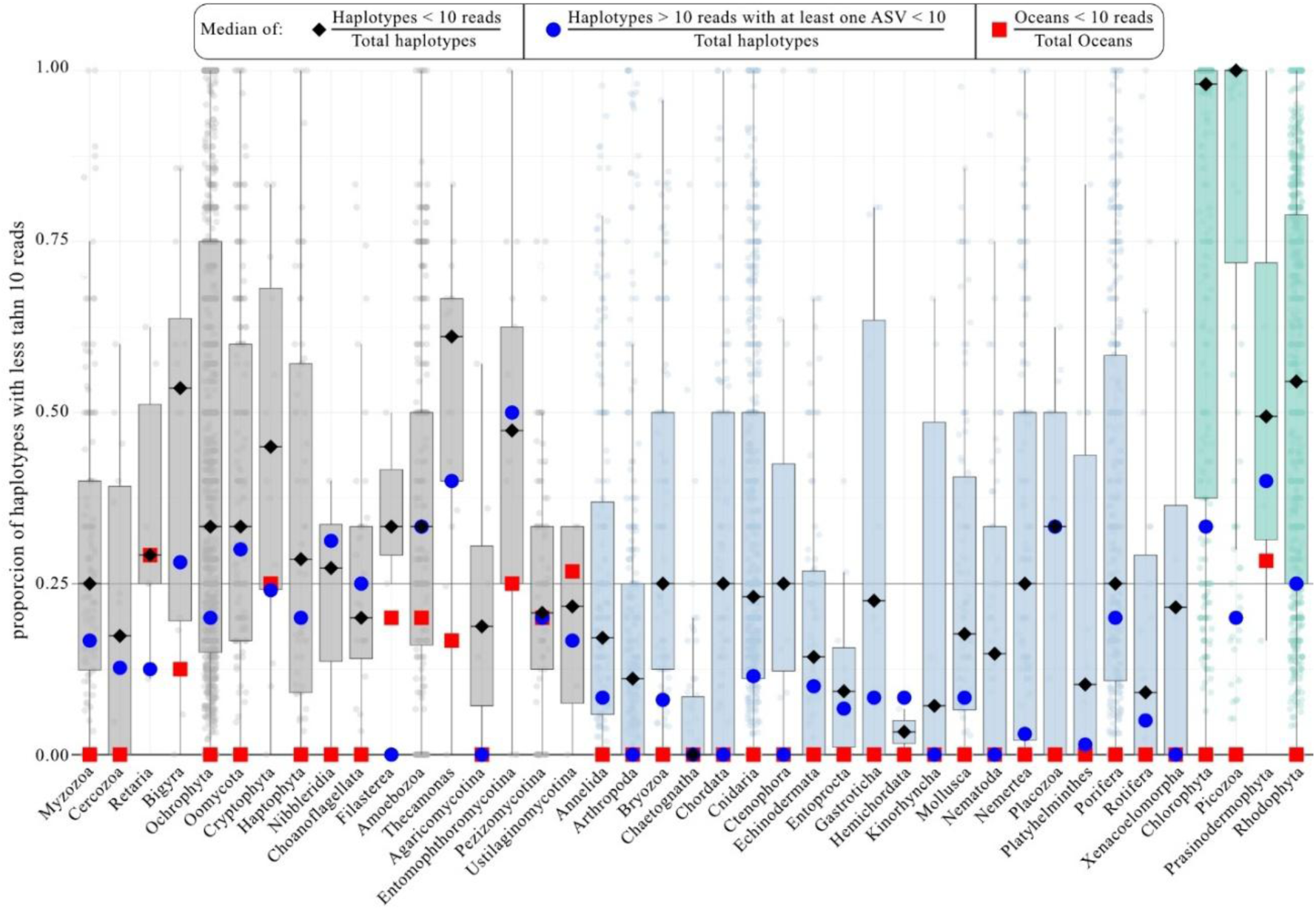
Rare haplotypes (< 10 reads). Boxplot represents the proportion of haplotypes with less than 10 total reads. The median is indicated by black diamonds, and the box colors correspond to different taxonomic groups. Blue circles represent the median proportion of haplotypes with more than 10 reads but composed of at least one ASV with less than 10 reads. Red squares represent the median proportion of oceans composed of less than 10 reads.

**Fig. S2.**
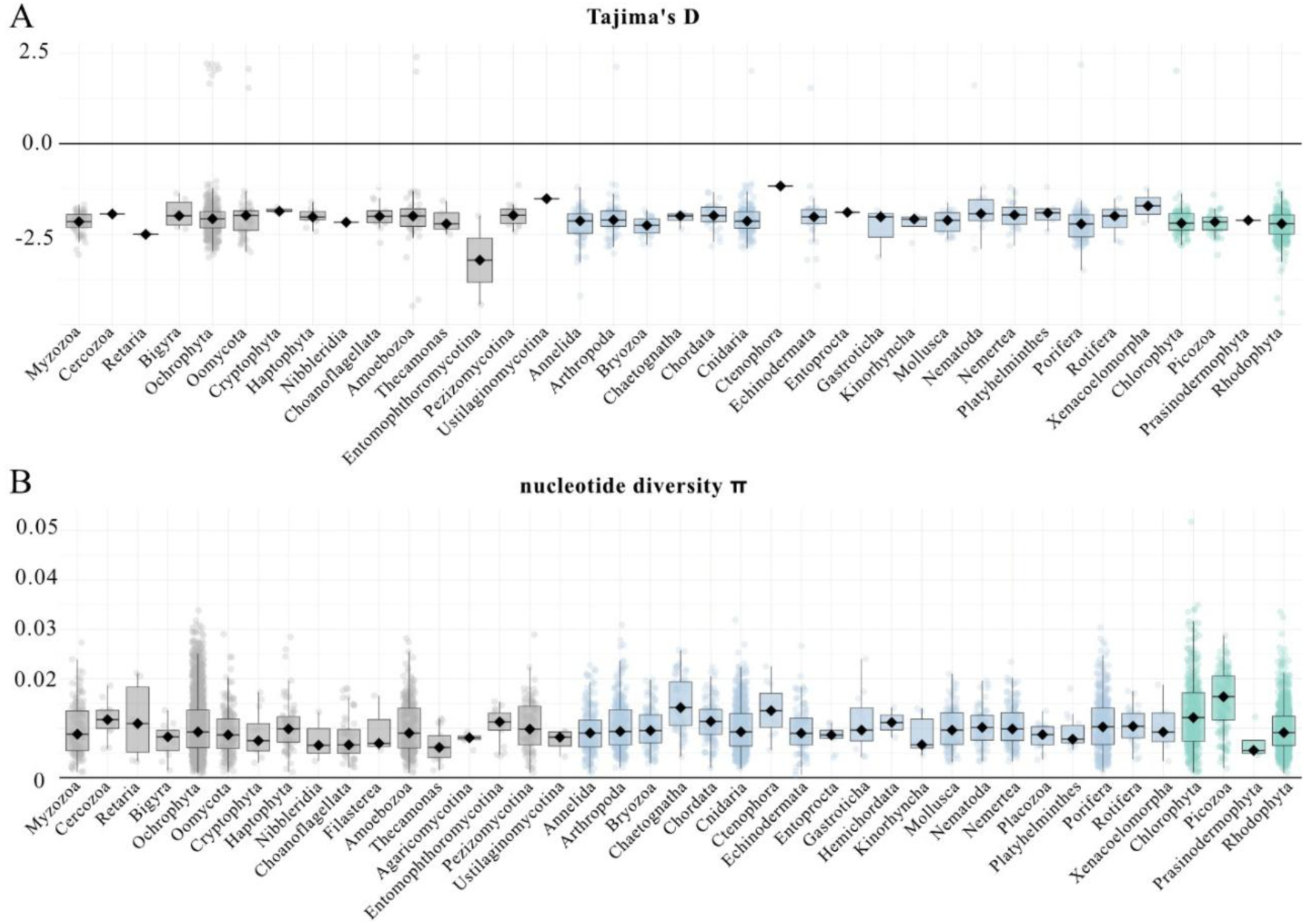
Nucleotide diversity metrics. (A) Boxplot representing the values of Tajima’s *D* for OTUs grouped by phyla, with p-values less than 0.05. (B) Boxplot of nucleotide diversity values for OTUs grouped by phyla.

**Fig. S3.**
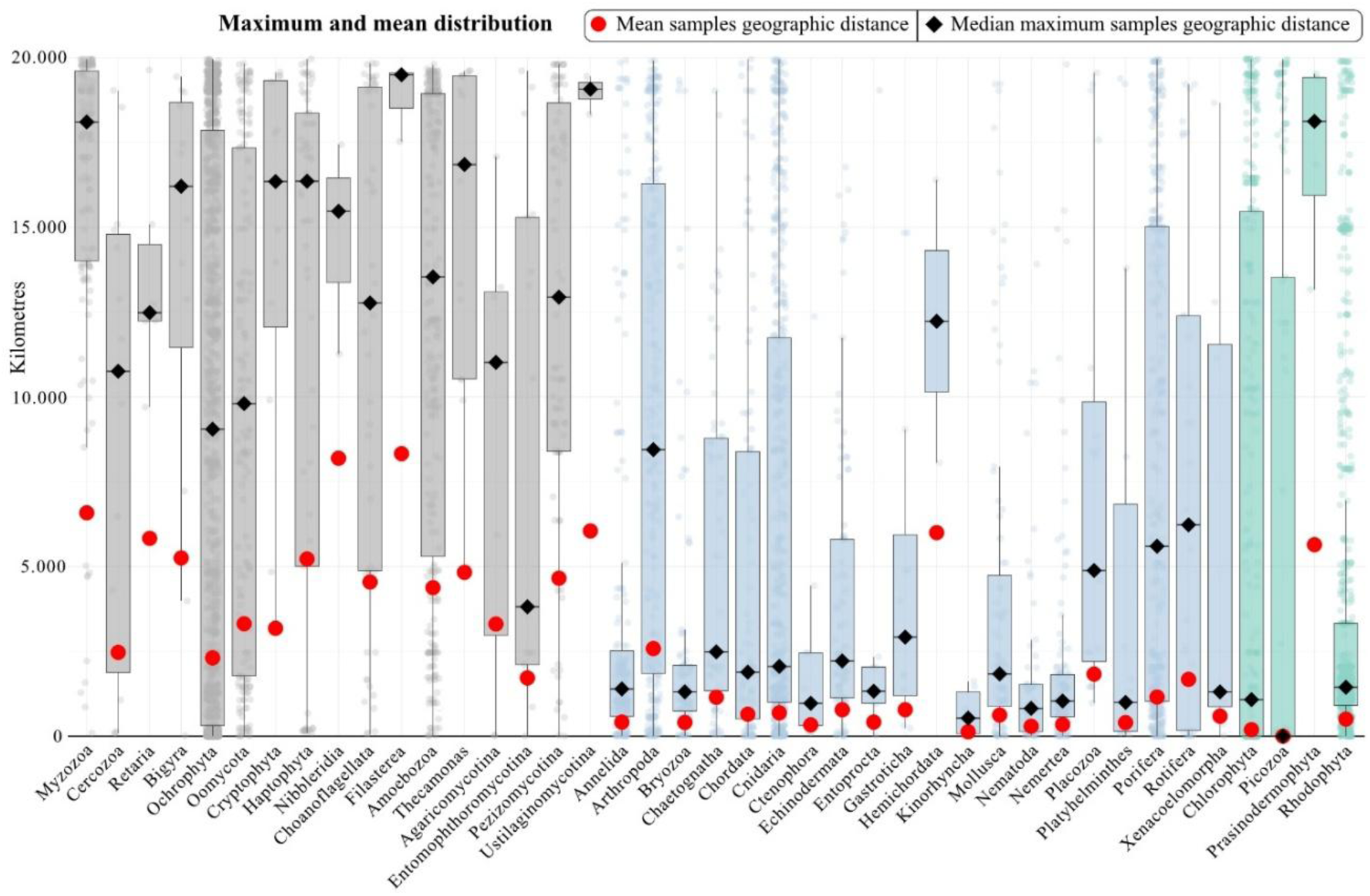
Geographic distances. The boxplots represents the maximum distribution between two localities per informative OTU. The red circles represents the median of the mean geographic distances between all the localities for each informative OTU.

**Fig. S4.**
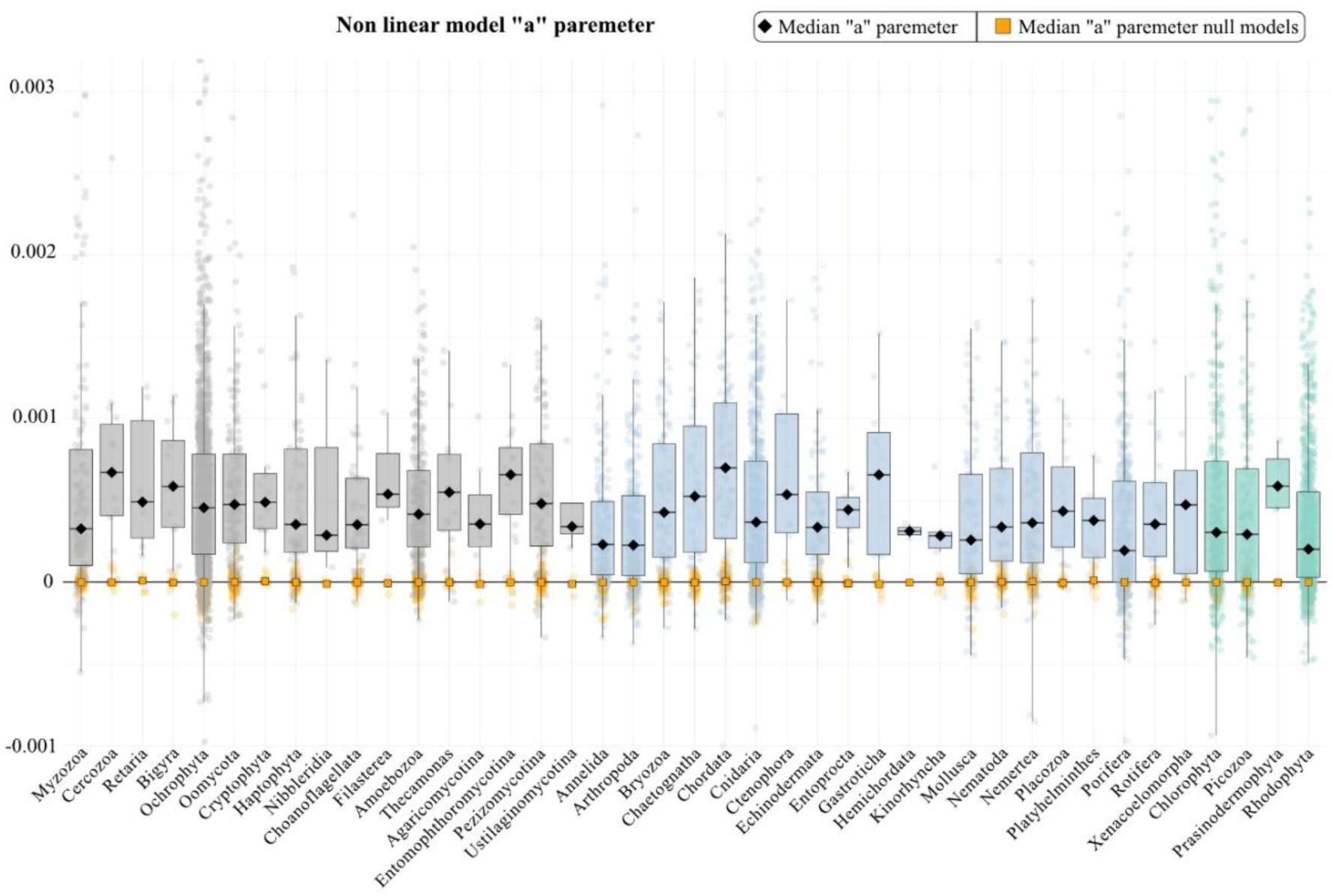
Parameter “a” of the nonlinear (exponential) model. Boxplots representing the values of parameter “a” from the non-linear models for each informative OTU, relating genetic and geographic distances. Orange points represent null models generated for each phyla, with squares marking the median values.

**Fig. S5.**
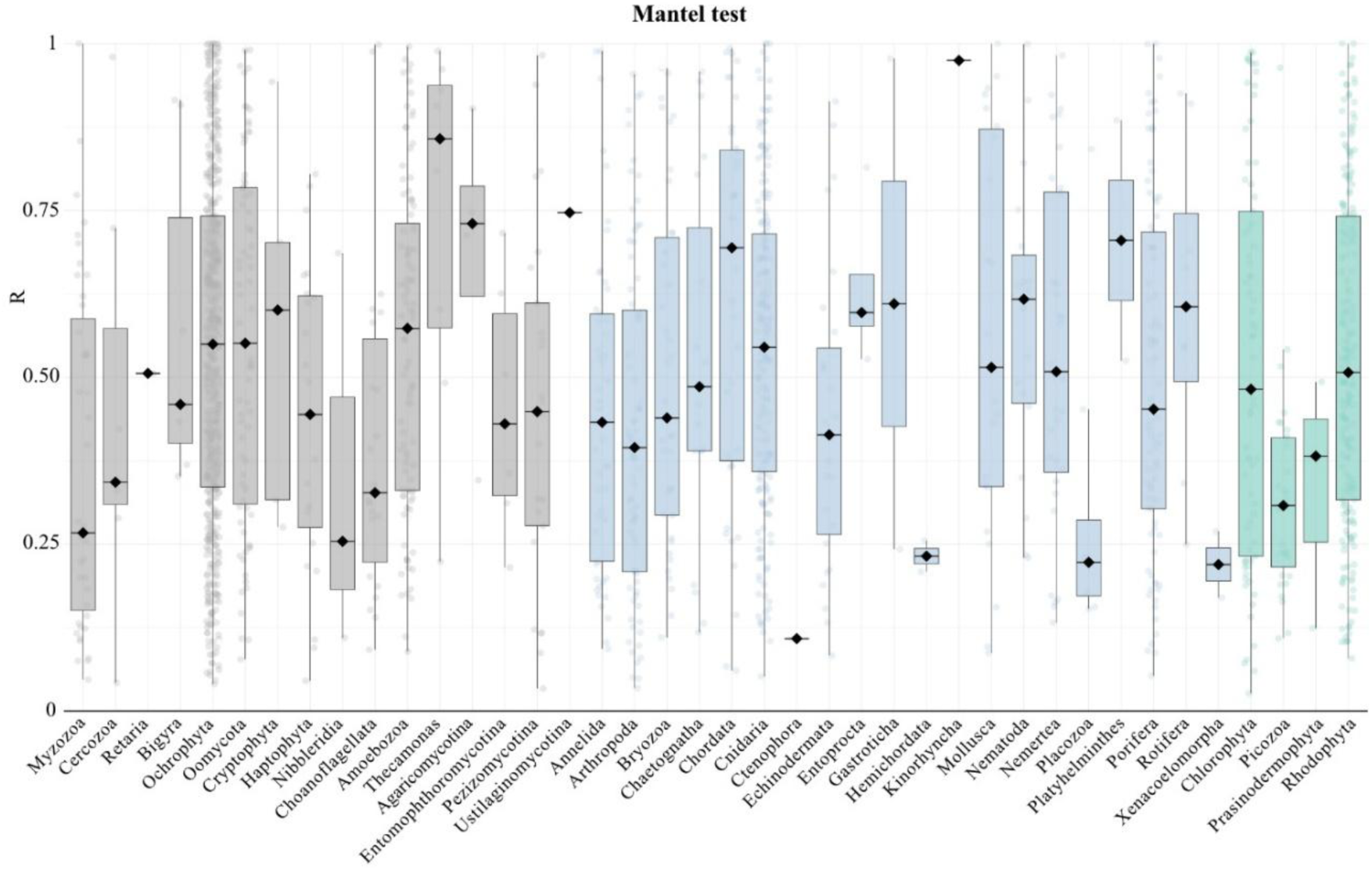
Significant Mantel test. Boxplots representing the *R* values from the Mantel test, including only informative OTUs with a p-value less than 0.05.

**Fig. S6.**
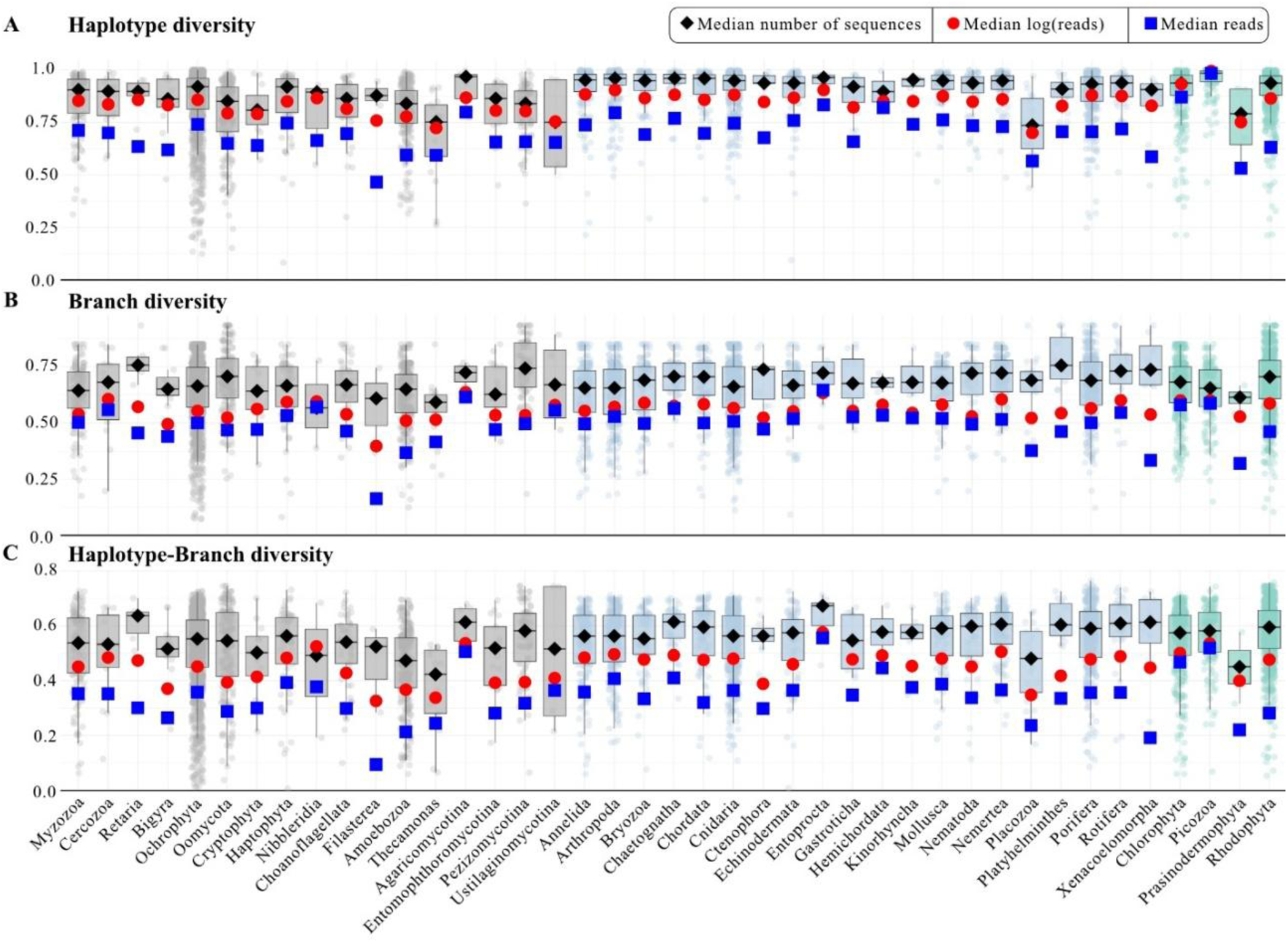
Haplotypic diversity metrics. (A) Boxplot representing haplotype diversity values, (B) branch diversity, and (C) haplotype-branch diversity, all based on the number of sequences. In all three cases, the median considering the number of sequences is represented by diamonds, the median based on total reads is represented by squares, and the median using log(reads) is represented by circles.

**Fig. S7.**
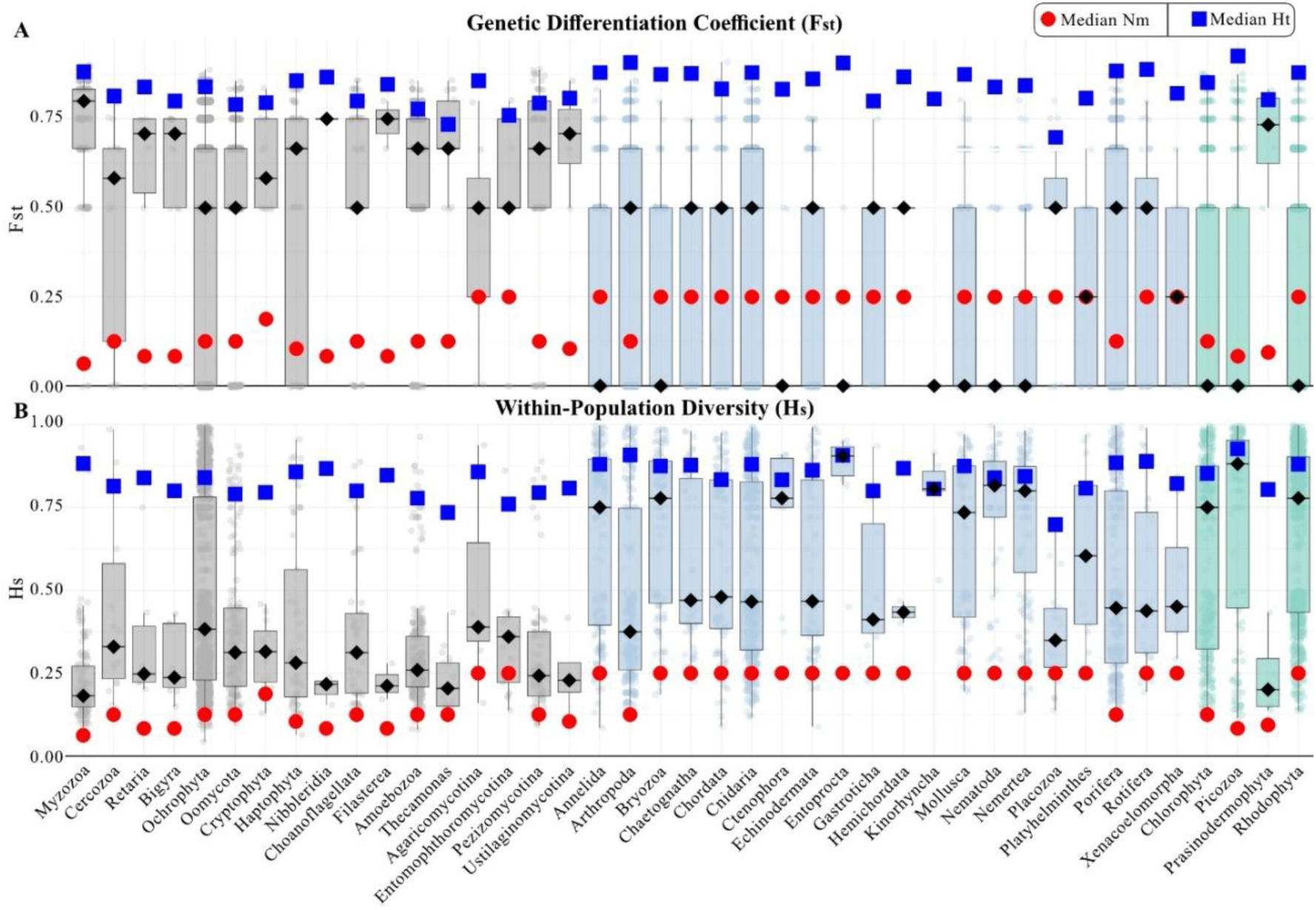
Population structure. (A) Boxplot representing the F_st_ values, and (B) boxplot of H_s_ values per informative OTU grouped by phyla. Circles represent the median of N_m_ values, and squares represent the medians for H_t_ values.

**Fig. S8.**
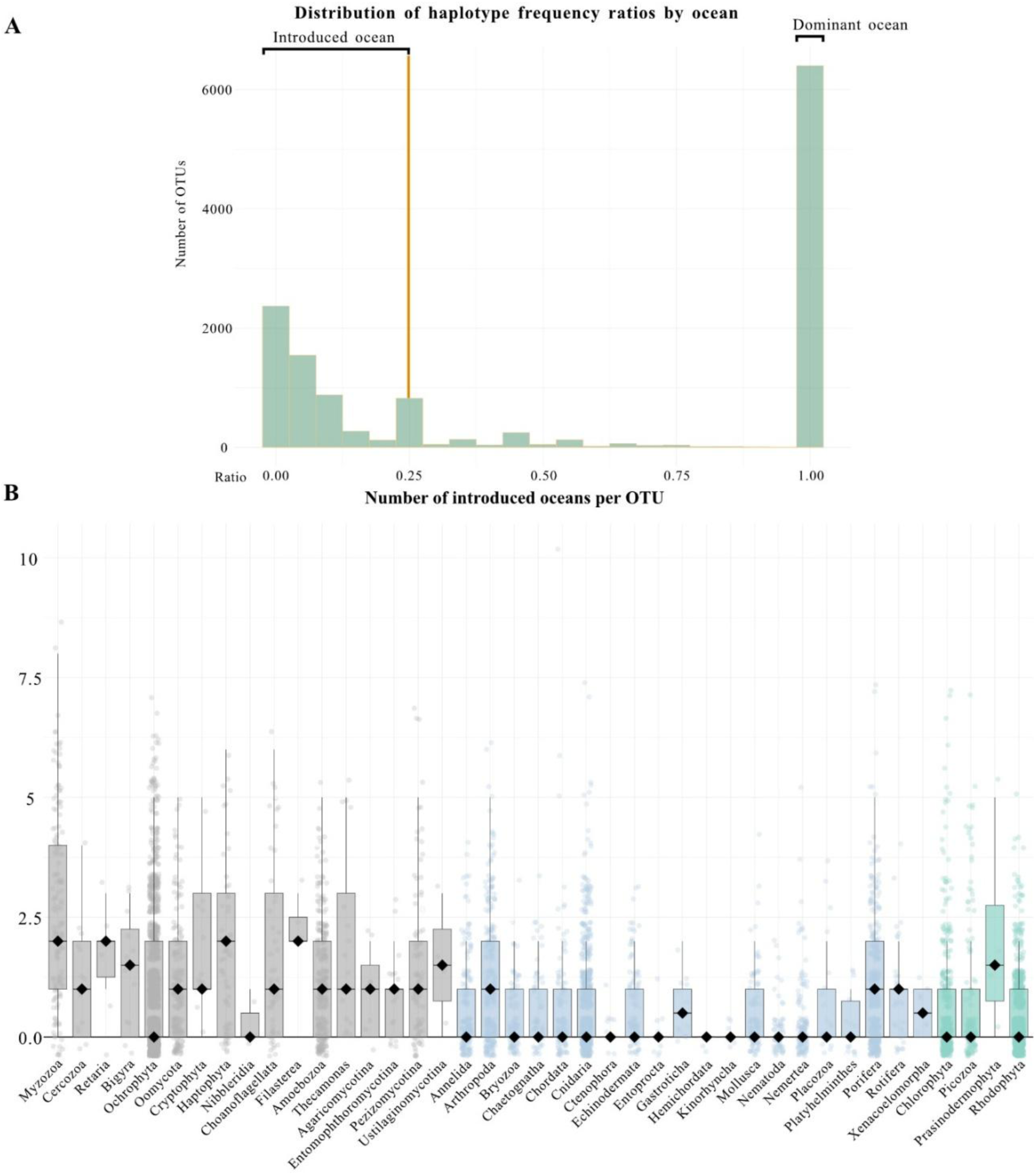
Characterization dominant and introduced ocean per OTU. (A) Distribution of haplotype relative frequency ratios between oceans of all informative OTUs. (B) number of introduced oceans per informative OTU grouped by phyla.

**Fig. S9.**
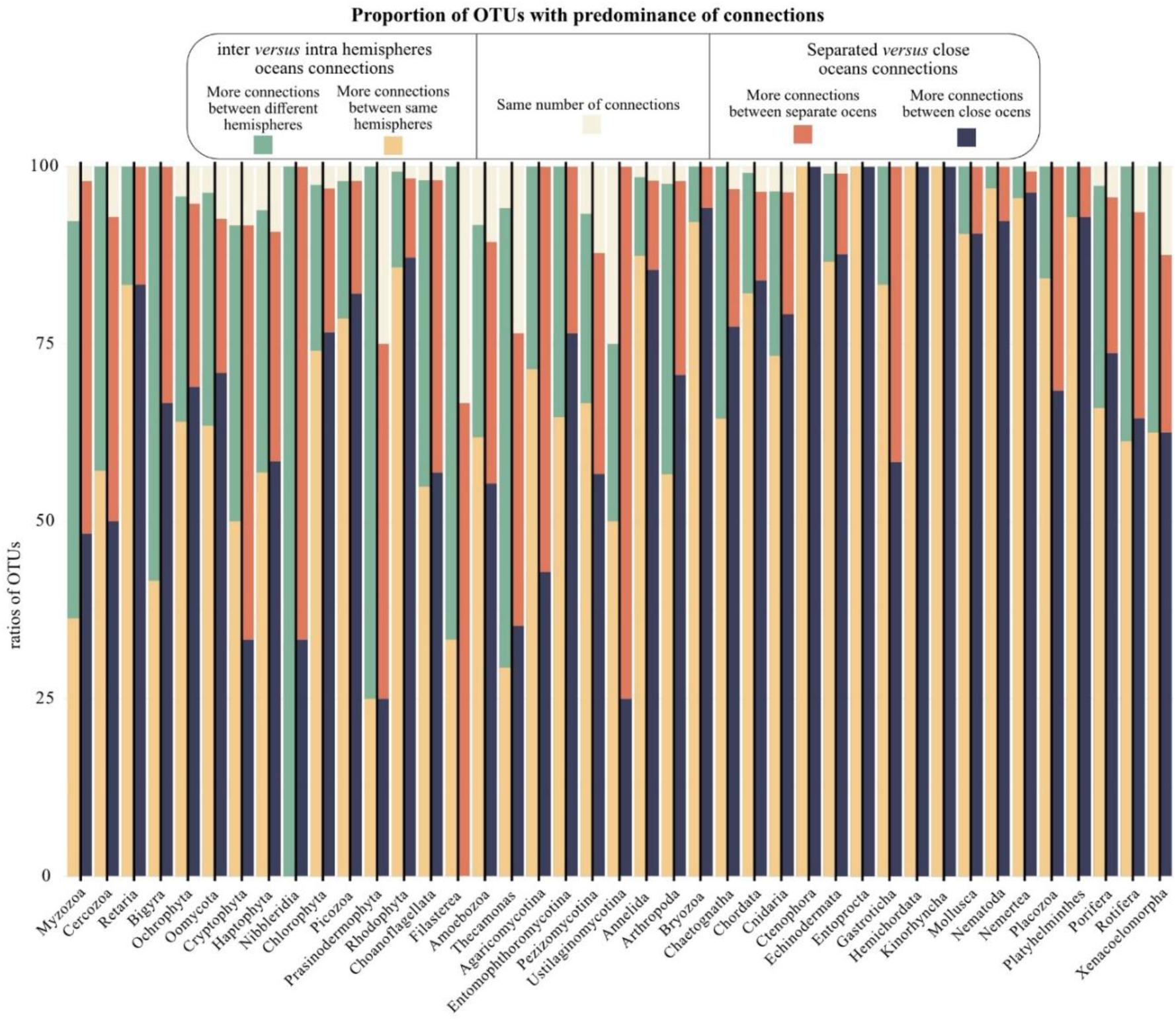
Ratios of haplotype oceanic connections. Bar plot representing the composition of the ratios for the different types of ocean connections, based on the haplotypic networks, for each phylum. The left side of the bar plot illustrates the proportion of connections between inter-versus intra-hemispheric oceans, while the right side shows the proportion of connections between separated versus adjacent oceans.

